# Phanta: Phage-inclusive profiling of human gut metagenomes

**DOI:** 10.1101/2022.08.05.502982

**Authors:** Yishay Pinto, Meenakshi Chakraborty, Navami Jain, Ami S Bhatt

**Author notes:** These authors contributed equally (Yishay Pinto, Meenakshi Chakraborty).

## Abstract

The human gut microbiome is a diverse ecosystem that encompasses multiple domains of life and plays a vital role in human health. Due to technical limitations, most microbiome studies have focused on gut prokaryotes, overlooking bacteriophages and other gut viruses. The most common method to profile viruses is to assemble shotgun metagenomic reads - often from virus-enriched samples - and identify viral genomes *de novo*. While valuable, this resource-intensive and reference-independent method has limited sensitivity. To overcome these drawbacks, we developed Phanta, which profiles human gut metagenomes in a virus-inclusive manner directly from short reads utilizing recently published catalogs of gut viral genomes. Phanta incorporates *k*-mer based classification tools and was developed with virus-specific properties in mind. Specifically, it includes optimizations considering viruses’ small genome size, sequence homology with prokaryotes, and interactions with other members of the gut microbial community. Based on simulations, the workflow is fast and accurate with respect to both prokaryotes and viruses, minimizing false positive species identification using a novel genome coverage-based strategy. When applied to metagenomes from healthy adults, Phanta identified ~200 viral species per sample, ~5x more than the standard assembly-based methods. Notably, we observed a 2:1 ratio between gut viruses and bacteria, with higher interindividual variability of the gut virome compared to the gut bacteriome. Phanta performs equally well on bulk vs. virus-enriched metagenomes, making it possible to study prokaryotes and viruses in a single experiment, with a single analysis. Phanta can tandemly profile gut viruses and prokaryotes in existing and novel datasets, and can therefore identify cross-domain interactions with likely relevance to human health. We expect that Phanta will reduce the barrier to virus-inclusive studies of the human gut microbiome, thus making it standard practice.

## Introduction

The human gut microbiome is an ecosystem of diverse microorganisms including archaea, bacteria, viruses, and fungi. It plays a vital role in human health by interacting with our immune, digestive, and nervous systems^1–4^. Since the 1970s, tools such as 16S rRNA sequencing have enabled us to identify prokaryotic taxa present in the gut^5^, and therefore to determine crucial relationships between these taxa and human health, age, lifestyle, environment, geography, and demographics^6–9^. However, these fundamental techniques overlook the viral fraction of the microbiome, preventing us from evaluating the impact of the human gut virome on human health.

Shotgun metagenomics is a popular and affordable method to sequence metagenomic samples^10–13^. This method captures genomic DNA from all gut organisms, not only prokaryotes, making it an optimal tool to study DNA viruses of the virome^14–16^. In the past decade, thousands of human microbiome samples have been analyzed using this “domain-inclusive” method^17–20^. Human gut prokaryotes can be well-quantified from shotgun metagenomes through direct read classification by comparing sequencing reads to reference genomes^18,19,21–24^. However, in the absence of comprehensive catalogs of viral genomes, the most common method for profiling the virome from shotgun metagenomes has been to assemble sequencing reads into contigs and identify viral genomes *de novo*^25,26^. Assembly-based approaches overcome the fundamental limitation that, until recently, a majority of phages had no reference genome^27^. However, despite their strengths at *de novo* phage discovery, assembly-based approaches have limited ability to detect low-abundance phages, due to the relative difficulty of assembling the genomes of low abundance taxa^28–30^.

With increases in shotgun metagenomes from human gut samples, more comprehensive databases of gut viral genomes have recently been created^27,31–37^. By using these new compendiums, it is now possible to profile gut viruses and their prokaryotic hosts simultaneously through read-based, reference-dependent methods. This approach can address the sensitivity limitation of assembly-based methods to profile the virome, resulting in much more complete profiles of the gut microbiome with both prokaryotes and viruses accurately represented.

In this paper, we present Phanta, a fast and accurate virus-inclusive profiler of human gut metagenomes based on classification of short reads to our newly constructed, comprehensive database of human gut microbes. The provided database contains the latest genome catalogs from multiple domains of life, including more than 190,000 phage genomes and the entire HumGut collection of prokaryotic genomes^19,27^. Phanta incorporates the state-of-the-art tools Kraken2^22^ and Bracken^38^, and complements them with additional filtering steps and optimizations specifically tailored to the challenges of gut viral quantification. Phanta accurately quantifies both bacteria and phage abundances in simulated mixed communities. In metagenomes from healthy human adults, Phanta identifies >100-fold more viral reads and minimizes unclassified reads when compared to the default Kraken2/Bracken databases and workflow. In addition, due to its high sensitivity, Phanta identifies 5-fold more viral species than a common workflow of contig assembly and viral sequence identification. Finally, Phanta quantifies just as many viruses when applied to bulk shotgun metagenomes vs. matched metagenomes enriched for virus-like particles. This demonstrates that it is possible to profile multiple domains of life from a single metagenomic sequencing experiment, as opposed to needing an additional sequencing experiment after enrichment for virus-like particles. Taken together, we anticipate that Phanta, which is freely available at https://github.com/bhattlab/phanta, will facilitate improved profiling of cross-domain interactions in gut microbiomes.

## Results

### Phanta: A workflow for phage-inclusive profiling of human gut metagenomes

Phanta was developed to generate accurate and complete profiles of human gut metagenomes, with the goal of deepening our understanding of cross-domain interactions in the gut. To achieve this objective, we first constructed a comprehensive database of gut microbial genomes found in humans. To minimize false mapping, it was important to curate comprehensive collections of genomes from all groups of taxa residing in the human gut - not only phages and other viruses, but also prokaryotes, eukaryotes, and possible contaminants. For this purpose, we used the HumGut collection as a reference for both human gut bacteria and archaea^19^. HumGut includes dereplicated genomes from both UHGG and RefSeq. For viruses, we used the Metagenomic Gut Virus catalog (MGV; dominated by human gut phages)^21,27^ and RefSeq. For gut eukaryotes, we also used RefSeq, and for contaminants, we used the human genome (hg38) and the Core UniVec database from NCBI^22^. To create an informative viral taxonomy, MGV genomes were first clustered to species-level operational taxonomic units (vOTUs). MGV vOTUs with high similarity to a RefSeq viral species were labeled with the NCBI-assigned taxonomy of that species. For the remaining MGV vOTUs, higher levels of taxonomy were assigned iteratively (see Methods).

The first step of Phanta is read classification to a database of reference genomes, such as that described above (Fig. 1A). As viruses have relatively low abundance in a typical metagenomic sample, we chose to use whole genome classification, which is typically more sensitive in the low-coverage regime than methods relying on clade-specific marker genes^22,39^. Specifically, Phanta classifies reads to the lowest possible taxonomic rank by Kraken2^22,24^, a *k*-mer-based method that has been shown to be both fast and accurate given the correct database and optimized parameters^39^. Second, Phanta reduces false positive species by filtering out species based on a calculated proxy for genome coverage (see Methods), a known issue in taxonomic classification^40^. Third, Phanta quantifies species-level relative abundances by executing Bracken, a tool complementary to Kraken2 that redistributes all classified reads to the species level using a Bayesian inference approach^38^. By default, Bracken calculates the “relative read abundance” - the proportion of reads assigned to a species out of all reads. However, since viral genomes can be orders of magnitude smaller than prokaryotic genomes, read abundance approaches inflate the relative signal from prokaryotes within a community. Therefore, we additionally calculate “relative taxonomic abundance”, which instead estimates the relative proportion of different organisms (not proportion of DNA sequence) within a given sample ^41^. Briefly, we adjust the relative read abundance of each species using the median length of the species’ genomes. This provides a comparable abundance estimation to amplicon sequencing or marker gene-based approaches (Fig. 1B). Lastly, Phanta allows users to determine cross-domain relationships by summing viral abundances by predicted host, providing information about the predicted virulence of the viral community, and correlating the abundances of phages and bacteria.

**Figure 1.**
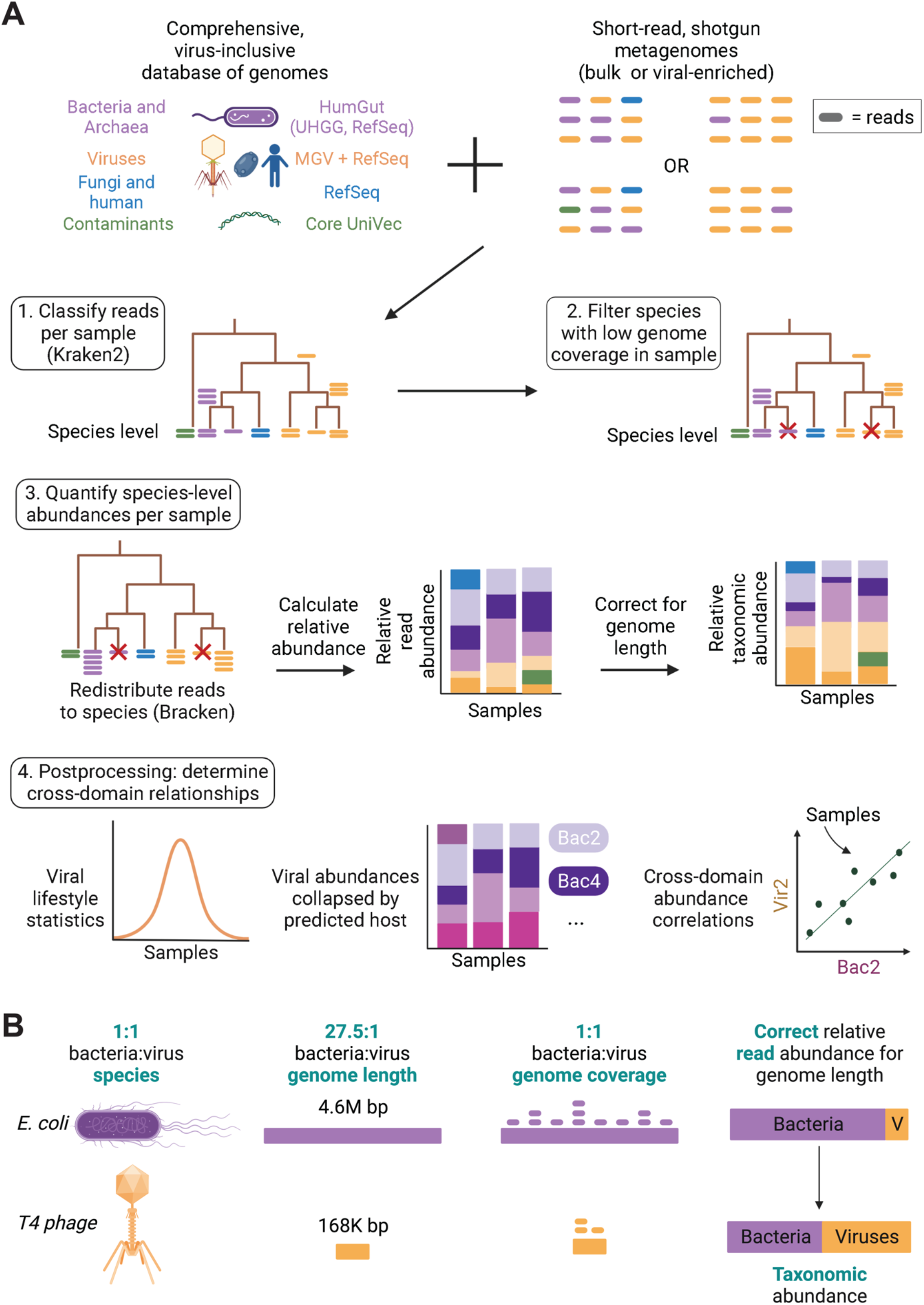
Overview of Phanta’s comprehensive, virus-inclusive metagenomic annotation workflow. **(A)** Phanta’s workflow. First, reads from each sample are classified against a comprehensive, virus-inclusive database of genomes from the human gut. Reads are classified to the lowest possible taxonomic level. After classification, genome coverage is estimated for each detected species in each sample. Species with low estimated genome coverage are filtered out to prevent false positive identifications. Next, Phanta quantifies the abundances of the remaining species in each sample. Reads originally classified above the species level (for example to the genus or family level) are redistributed downwards. Then, two types of abundance are calculated: (1) relative read abundance, which normalizes species-level read counts by read depth, and (2) relative taxonomic abundance (see panel B). Post-processing scripts are provided to determine cross-domain relationships. **(B)** Motivation behind Phanta’s provided correction of relative read abundance of relative taxonomic abundance. Shown here is a simple gut microbial community with a 1:1 ratio between bacteria and viruses (one bacterium of species *E. coli;* one virus of species T4). Even if *E. coli* and T4 genomes are equally covered by reads in a shotgun metagenome, the dramatic difference between their genome lengths will inflate the ratio of bacteria to viruses, if relative read abundance is used as the metric. By contrast, relative taxonomic abundance, which corrects for genome length, accurately captures the 1:1 ratio of these species.

### Phanta accurately classifies short reads from simulated mixed microbial communities

To evaluate the performance of Phanta, we simulated 10 mixed communities, each containing a total of ~6.5M 150 base pair (bp) paired-end reads from a combination of 300 prokaryotic genomes and 50 viral genomes (see Methods). The relative read abundance of prokaryotes and viruses in the resulting simulated samples was 0.95 and 0.05, respectively (Figure 2A). Phanta accurately assigned reads to the right domain with average read abundance of 0.951±0.004 mean read abundance for prokaryotes, and 0.048±0.004 mean read abundance for viruses (Figure 2B; Supplementary Data File 1).

**Figure 2.**
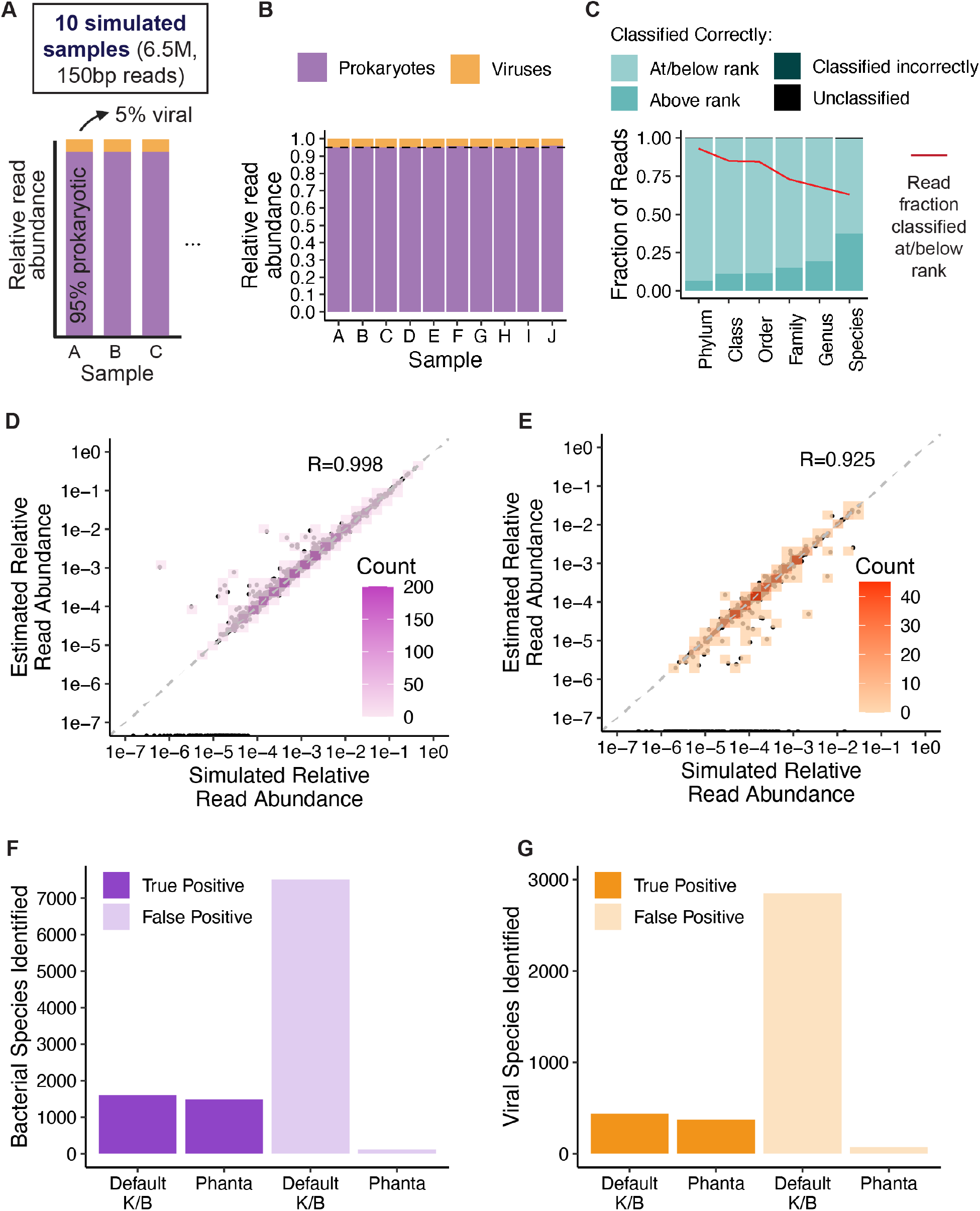
Evaluation of Phanta’s performance using simulated metagenomes. **(A)** Composition of simulated metagenomes. Results in (B) - (G) were obtained by applying Phanta to these simulated metagenomes while using Phanta’s default database and parameters. **(B)** Accuracy of Phanta’s final estimates of relative read abundance at the domain level. The dashed line indicates the true relative read abundance of prokaryotes. **(C)** Accuracy of Phanta’s classification step at each taxonomic rank. For each rank, the two shades of blue represent reads that were classified to a lineage that included the correct value of the rank. Specifically, light blue shading indicates the median fraction of reads (across simulated samples) that were classified correctly at or below the rank - e.g. for family, they were classified either to the correct family, or to the correct genus/species, which is even more specific than the correct family. Darker blue shading indicates the median fraction of reads that were classified correctly above the rank - e.g. for family, they were classified to the correct order or phylum, which is less specific than the correct family but still accurate. The dark green and black portions of each bar represent reads that were either: (i) classified to a lineage that did not include the correct value of the rank, or (ii) unclassified, respectively. The red line indicates the median fraction of classified reads that were classified at or below each rank (e.g., what fraction of reads were classified at or below the family level). **(D)** Accuracy of Phanta’s final estimates of relative read abundance for 1,606 bacterial species, across all simulated samples. The dashed line is the x=y diagonal. Each dot represents one bacterial species in one simulated sample. The x-axis is the simulated abundance in the sample, and the y-axis is the abundance estimated by Phanta. The *R* value indicates Pearson’s correlation coefficient, considering all the dots, i.e. all bacterial species in all simulated samples. Colors of the overlaid boxes represent numbers (counts) of dots. **(E)** Same as (D), for 500 viral species. **(F)** Signal-to-noise ratio of bacterial species identified by Phanta vs. the Kraken2/Bracken workflow, using default parameters for both workflows and using Phanta’s default database as the reference database. **(G)** Same as (F), for viral species.

We next used the simulated communities to test the accuracy of classification of reads by Kraken2. Reads were classified with high precision to all taxonomic ranks, with 63% of reads classified to the species level or lower (median across simulated communities; see Figure 2C). Next, we tested the accuracy of Phanta in estimating the abundance of each simulated species. Phanta’s species-level estimates for relative read abundance were highly correlated with the true simulated values - Pearson’s R=0.997 for all species (including bacteria and archaea), R=0.998 for bacterial species (Figure 2D), and R=0.925 for viral species (Figure 2E).

### Phanta’s filtering step significantly reduces false positive species identification

While developing Phanta, we observed that even a small fraction of mis-classified reads can lead to a non-negligible number of falsely identified species. Therefore, to increase the signal-to-noise of the identified species, we made the following modifications to the default Kraken2-Bracken workflow. First, we introduced a filtering step between Kraken2 and Bracken that estimates the breadth of genome coverage for species detected by Kraken2 and filters out likely false positive species based on a user-adjustable coverage threshold. In addition, for a read to be classified by Kraken2, we required that a certain fraction of a read’s *k*-mers be mapped to a given taxon, in order for the read to receive that classification. To achieve this, we adjusted Kraken2’s confidence threshold. By default, Phanta uses a confidence threshold of 0.1 (vs. 0 for default Kraken2; also recommended by ^39^), and this can be further adjusted by the user. These steps reduced false positive species by 50-fold with minimal reduction of true positive species relative to a consecutive run of Kraken2 and Bracken using default parameters (Figures 2F-G). Overall, we demonstrated that Phanta performs with high accuracy in both classifying reads and estimating abundance while substantially reducing false species identification.

### Masking prophages in prokaryotic genomes further increases sensitivity to viral reads

Due to genetic flow between viruses and their hosts, phage genomes share a relatively high proportion of their genome with their bacterial hosts (Supplementary Fig. 1A). This can limit detection of viral sequences in metagenomes, because portions of the viral sequences will also be present in bacterial genomes. Therefore, we decided to construct an alternative version of Phanta’s default database, in which prophage sequences, which are phage sequences that are integrated into the bacterial chromosome, are “masked”. This is accomplished by replacing the prophage sequences with Ns in all bacterial genomes where they appear. Prophage sequences were predicted using VIBRANT^42^. We anticipated that masking would further increase Phanta’s sensitivity to viral reads in simulated communities. Indeed, using the masked database reduced the number of “ambiguous” read classifications - i.e., the number of reads that Kraken2 classified to the “root” of the taxonomy tree. The vast majority of reads that were classified to the root using the default database, but received a new classification after masking, were reclassified to the viral domain (Supplementary Fig. 1B). This result demonstrates that: (1) shared sequences between bacteria and viruses can indeed result in ambiguous read classification, and (2) this ambiguity can be partially resolved by masking prophages in bacterial genomes. Importantly, masking does not lead to over-detection of viruses; Phanta’s final read abundance estimate for viruses remained highly accurate (Supplementary Fig. 1C).

### Phanta improves the overall proportion of reads classified in shotgun metagenomes from healthy adults

Given the good performance of Phanta on simulated samples, we wished to assess whether Phanta could improve viral identification in samples from healthy adults. We applied Phanta with the default (no prophage masking) database on human gut metagenomes sampled from 245 healthy adults (age range 21-79, from Yachida *et al*.)^43^. In total, across 245 samples, the workflow took ~60 minutes to run using 1 core, 16 threads, and 32GB memory. Given that Phanta incorporates Kraken2 and Bracken, we were easily able to compare the workflow’s performance using Phanta’s default database, compared to existing Kraken2/Bracken-compatible databases. In particular, we compared against four existing databases (Table 1): the standard Kraken2 database^22,44^ (May 2021), the Unified Human Gastrointestinal Genome (UHGG) collection^18^ (July 2021), RefSeq Complete^39^ (April 2022), and HumGut^19^ (July 2021). Phanta’s default database was able to minimize the number of unclassified reads to 2% (Fig. 3A), and notably, it requires ~97% less disk space than the most comprehensive database tested, RefSeq Complete (32GB for Phanta, 1.2 TB for RefSeq Complete^39^).

**Table 1.** Characteristics of the different Kraken2/Bracken-compatible databases tested in this study.

| Database | Size (GB) | Prokaryotic<br>genomes* | Viral<br>genomes* | Median<br>classification<br>time (sec)** | Median %<br>classified<br>reads** |
| --- | --- | --- | --- | --- | --- |
| Standard<br>Kraken2 | 51 | 21,920 | 10,489 | 491 | 56.31 |
| RefSeq<br>Complete | 1,192 | 215,725 | 10,863 | 4,889 | 93.61 |
| UHGG | 16 | 4,644 | 0 | 548 | 88.75 |
| HumGut | 26 | 30,691 | 0 | 549 | 97.57 |
| Phanta | 32 | 30,691 | 201,305 | 544 | 98.07 |

\*Numbers were obtained from the original papers, see Methods. \*\*Classification times and percentages of classified reads were determined by conducting Kraken2 classification of five random samples from the healthy human cohort from Figure 3, and calculating median classification times and percentages across the samples.

**Figure 3.**
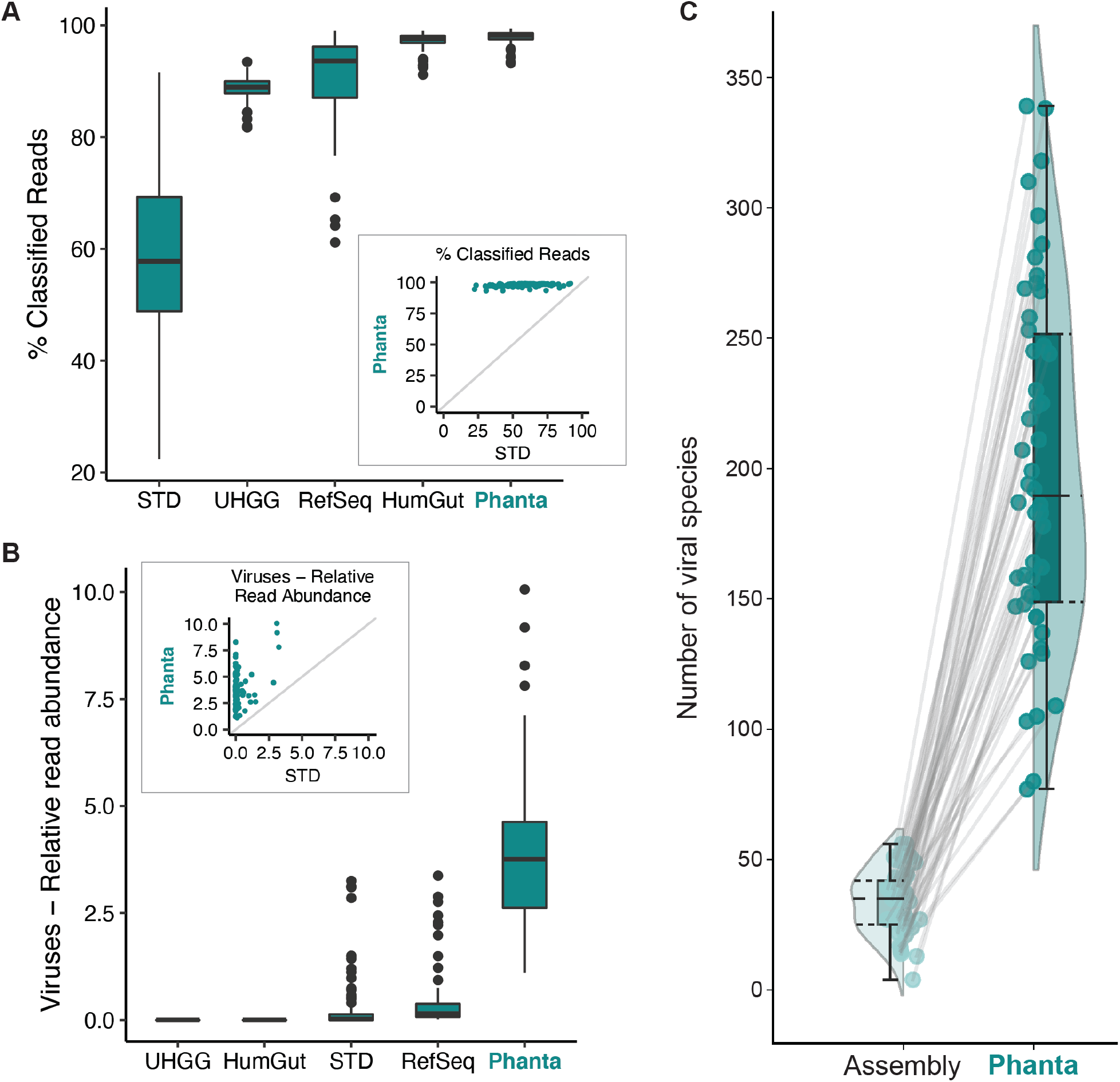
Evaluation of Phanta’s performance using shotgun gut metagenomes from 245 healthy human adults. Metagenomes sourced from Yachida *et al*.^43^ **(A)** Percentage of sample reads that could be classified during Phanta’s initial classification step using Phanta’s default parameters and a variety of Kraken2/Bracken-compatible databases. Boxplots display the percentage distribution across the set of metagenomes. Database abbreviations: STD = standard Kraken2^44^, UHGG = Unified Human Gastrointestinal Genome Collection^18^, RefSeq = RefSeq Complete v205^39^, HumGut = HumGut^19^, Phanta = Phanta’s default database. The insert shows the same information as the boxplots for STD and Phanta. **(B)** Similar to (A) but comparing the relative read abundance of viruses after Phanta’s filtering and abundance estimation steps. **(C)** Comparing the number of distinct viral species identified by Phanta using the default database and parameters vs. a standard, assembly-based workflow to identify viral species in shotgun metagenomes. Dots represent individual metagenomes and lines are drawn between dots representing the same metagenome. Distributions of dots are shown using both boxplots and violin plots. **Note:** in all boxplots, boxes represent the interquartile range (IQR), the horizontal line indicates the median, and whiskers extend between (25th percentile - 1.5*IQR) and (75th percentile + 1.5*IQR).

### Phanta substantially increases viral identification in shotgun metagenomes

In addition to maximizing classified reads, Phanta’s default database led to the highest level of viral identification, detecting 25-fold and 188-fold more viral sequences compared to RefSeq Complete and the standard Kraken2 database, respectively (Fig. 3B; Supplementary Data File 2). Using Phanta, we now estimate that viral DNA constitutes 3-5% of the DNA in the human gut. Taken together, Phanta improves read classification both by enabling the classification of previously unclassified reads and by improving the recognition of viral sequences.

### Phanta outperforms standard assembly-based methods in identifying viruses in shotgun metagenomes

A current gold-standard workflow commonly used to identify viruses in shotgun metagenomes involves assembling reads into contigs and labeling the likely viral contigs^42,45–49^. To compare Phanta to this gold standard, we randomly selected 50 metagenomes from the healthy adult cohort and ran a standard assembly workflow. In short, reads were assembled to contigs using metaSPAdes^50^, short/low-quality contigs were filtered using CheckV^51^, and viral contigs were identified using both VIBRANT^42^ and VirSorter^45^. For each sample, the total set of viral contigs from both methods was de-replicated to 95% ANI to calculate a number of viral species. Phanta was able to identify a higher number of viral species than the assembly workflow in all samples, with a median of 190 (IQR: 149-252) viral species per sample relative to 35 (IQR: 25-42) (Fig. 3C) identified with assembly-based approaches. Of note, the vast majority of viral contigs predicted by assembly were highly similar to genomes in the viral portion of Phanta’s database (Supplementary Fig. 2).

### There are twice as many viral as bacterial genomes in the human gut

By default, Bracken calculates relative read abundance for each identified taxon - i.e., the fraction of reads classified to it. This measurement serves as an estimation of the fraction of genomic DNA belonging to each taxon, out of the total DNA in a sample. While this measurement is highly valuable, an ecological perspective of a community requires understanding the proportions of “individuals” in the community - i.e., relative taxonomic abundance^41^. Relative read abundance is typically similar to taxonomic abundance in communities with similar genome lengths. However, in mixed communities containing taxa with orders of magnitude differences in genome length, like bacteria and viruses, relative read abundance is biased towards taxa with longer genomes (as illustrated in Fig. 1B). Hence, Phanta calculates an estimation of relative taxonomic abundance by correcting the relative read abundance by genome length. Using our relative taxonomic abundance calculation, we estimate the ratio between copies of viral genomes to bacterial genomes in the human gut to be ~2:1 (Figs. 4A-B; Supplementary Data File 2). Phanta also reports several other normalizations - reads per million base pairs, reads per million reads, reads per million base pairs per million reads (analogous to RPKM in transcriptomics) and genome copies per million reads.

**Figure 4.**
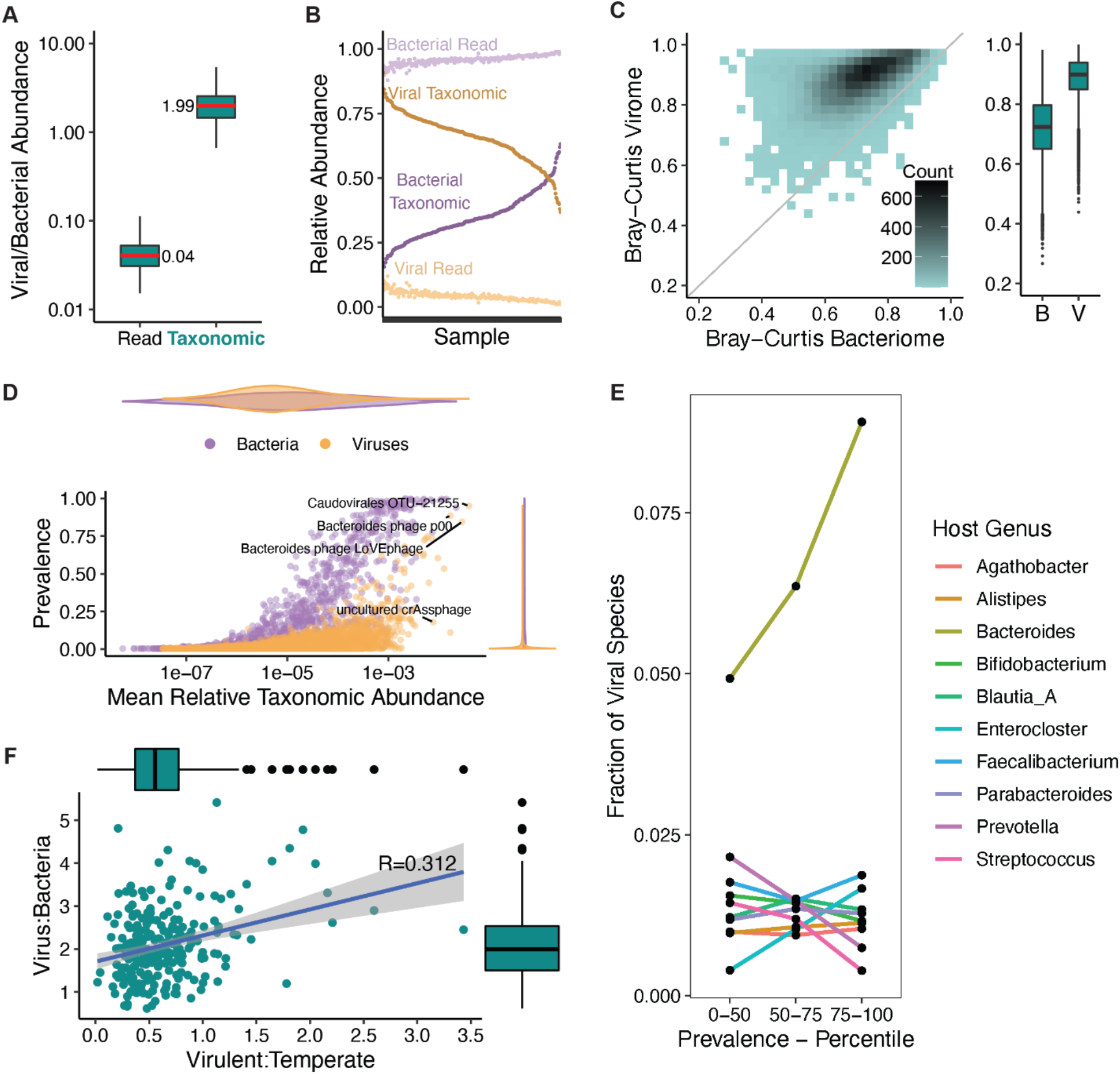
Core properties of the healthy adult virome. Metagenomes sourced from Yachida *et al*.^43^ (same as Figure 3). **(A)** Ratio of viral to bacterial abundance in the gut, using relative read abundance vs. relative taxonomic abundance. Boxplots display the distribution of this ratio across the set of 245 healthy adult metagenomes. **(B)** Abundance values used to calculate the ratios in (A). **(C)** Comparing the variability of the gut phageome and bacteriome across metagenomes. Bray-Curtis dissimilarities were calculated twice between all metagenome pairs, once using relative taxonomic abundances of bacterial species (horizontal axis of scatterplot) and once using relative taxonomic abundances of viral species (vertical axis of scatterplot). The boxplots display the same data - B = bacteriome, V = virome. The gray line on the scatterplot is the x=y diagonal. **(D)** Abundance and prevalence of bacterial and viral species. Abundance is the mean relative taxonomic abundance across metagenomes and prevalence is the number of positive individuals divided by the cohort size (245). Violin plots aligned with the x- and y-axes represent distributions of abundance and prevalence, respectively. **(E)** Distribution of predicted host genera for viral species in various prevalence categories (e.g., category 75-100 represents the top 25% of viruses in terms of prevalence). These results are based on host genus predictions that were made using iPHoP^65^ and are provided in Phanta’s default database. **(F)** Relationship between abundance ratio of viruses and bacteria and abundance ratio of virulent and temperate phages. Boxplots aligned with the x- and y-axes display the distributions of each ratio. Results are based on viral lifestyle predictions made by BACPHLIP^64^ (provided in Phanta’s default database). Displayed *R* is Pearson’s correlation coefficient. Relative taxonomic abundance was used as the abundance metric. To prevent low quality samples from affecting the analysis, 11 outliers for sequencing depth - i.e., >1.5*IQR above or below the median depth - were removed (n=234). **Note:** in all boxplots, boxes represent the interquartile range (IQR), the horizontal line indicates the median, and whiskers extend between (25th percentile - 1.5*IQR) and (75th percentile + 1.5*IQR).

### High individuality of the human gut virome

We further used the viral and bacterial profiles reported by Phanta to describe core differences between the virome and bacteriome of healthy adults. We observed a higher between-sample dissimilarity of the virome relative to the bacteriome in healthy adults (Fig. 4C). The high dissimilarity of the virome between individuals points to a highly personalized virome, as has been suggested previously^31,52–54^. Consistent with this result, individual viral species are skewed towards lower prevalence than bacterial species (Fig. 4D). However, a number of lowly prevalent viruses show high mean abundance across individuals, indicating that they are highly abundant when present. As previously suggested, the prototypical crAssphage^55,56^ (RefSeq ID 1211417) was one of the most abundant viral species, although it was not among the most prevalent (Fig. 4D and Supplementary Table 1). Two of the most prevalent and abundant species were OTU-66229 and OTU-72541. These phages are highly similar to the recently described Bacteroides phages LoVEphage^37^ and Hankyphage (p00)^57^, respectively (Supplementary Fig. 3). The most abundant and prevalent phage detected was Caudovirales OTU-21255, a temperate phage likely of family Siphoviridae whose presumed host is *Bacteroides uniformis*. This species was found in 232/245 (95%) of individuals in this cohort of healthy adults, and comprises 1512 genomes in Phanta’s default database.

### Prevalent phages infect Bacteroides

We next examined relationships between viral species prevalence and predicted host. Bacteroides is the most commonly predicted host genus for viral species detected in the healthy adult cohort. Specifically, it was the predicted host for 6.5% of detected viral species, compared with 3% of species in Phanta’s database, more than twice than expected. The dominance of Bacteroides as a predicted host further increases among the more prevalent viral species (Fig. 4E).

### Temperate phages dominate the human gut phageome

Phanta’s default database includes estimates of virulence per species (see Methods), which we used to determine the ratio between different phage lifestyles (virulent vs. temperate) in the human gut. We observed that in the vast majority of samples temperate phages are dominant with a median of 0.54 for the ratio of virulent/temperate species identified, and 0.55 for the corresponding abundance ratio. Notably, more prevalent phages are skewed towards the temperate lifestyle (Supplementary Fig. 4), potentially reflecting the ability of some temperate phages to remain dormant in their hosts. Interestingly, the abundance of virulent phages in the community, relative to temperate phages, is positively correlated with overall phage abundance in the microbiome (Fig. 4F). This is consistent with the nature of virulent phages, whose active replication increases their ratio relative to their bacterial host.

### Phanta performs well on virus-enriched metagenomes

Viral enrichment, either through filtration or other approaches to achieve viral particle isolation, is commonly used in viromics studies to enhance the detection of viral DNA in metagenomes^58^. Therefore, we wanted to test Phanta’s performance in metagenomes originating from virus-enriched samples. We applied Phanta to paired bulk and virus-enriched shotgun metagenomes from infants (Supplementary Data Files 3-6; source data: Liang *et al*.^59^). We first tested the performance of Phanta on the virus-enriched samples by correlating the viral-like particle counts (from Supplementary Table 2 in ^59^) to the number of viral species identified (i.e., viral species richness) by various assembly or classification methods. Phanta-based richness was the most strongly correlated with VLP counts (Fig. 5A).

**Figure 5.**
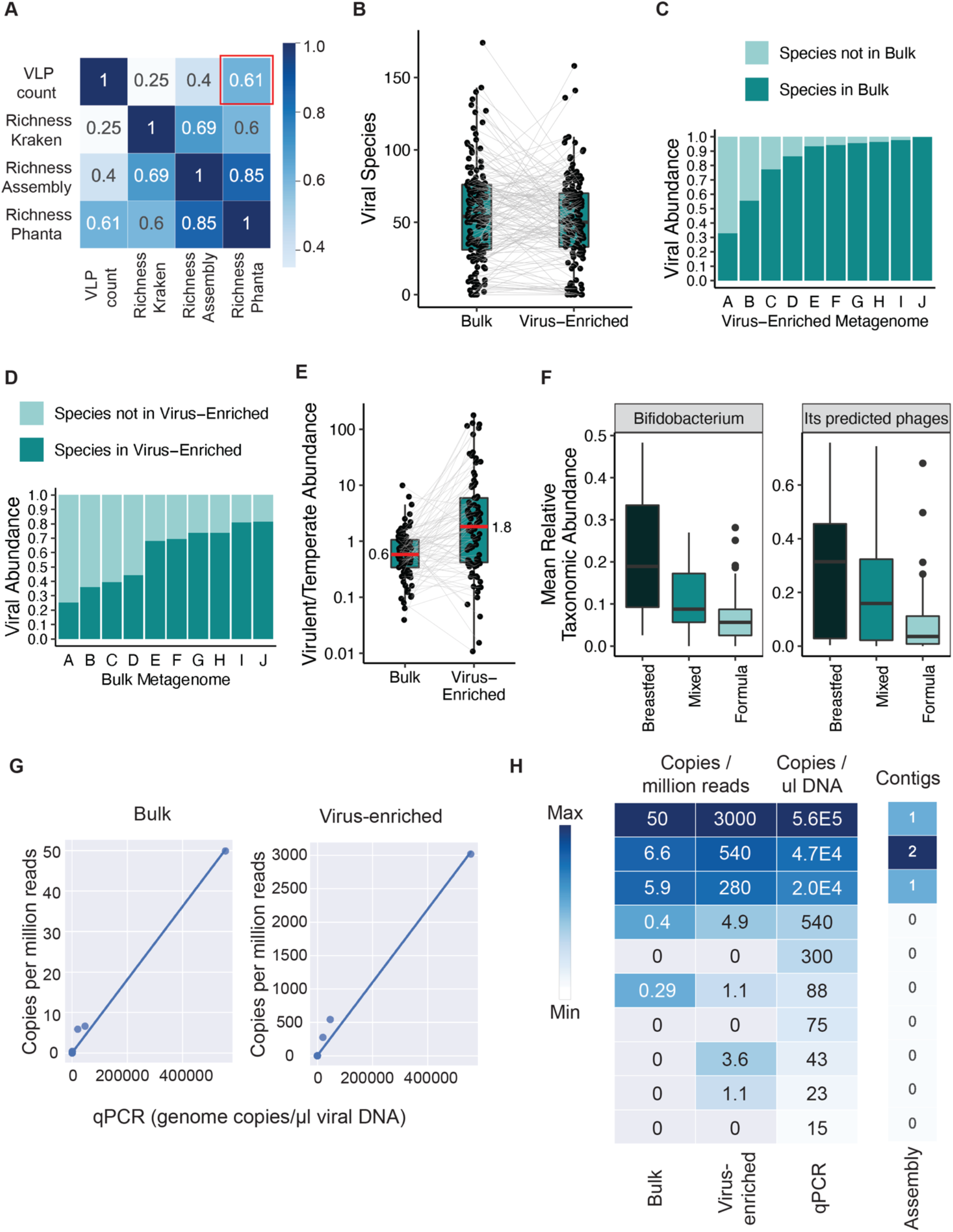
Application of Phanta to paired virus-enriched and bulk metagenomes from the infant gut. Metagenomes sourced from Liang *et al*.^59^ Longitudinal cohort = 20 infants sampled at months 0, 1, and 4 (60 samples total). Four-month cohort = 83 infants sampled at month 4. **(A)** All-by-all Spearman’s correlations between statistics related to viral content, for all virus-enriched metagenomes from infants in the longitudinal cohort (n=60). Specifically, four statistics were correlated: (1) VLP Count: number of viral-like particles per gram feces, (2) Richness Kraken: viral species richness based on applying Kraken2 with a RefSeq-based database, (3) Richness Assembly: viral species richness based on applying an assembly-based method, and (4) Richness Phanta: viral species richness based on applying Phanta. Phanta’s richness estimation has the highest correlation with VLP count (red box). VLP Count, Richness Kraken and Richness Assembly were originally reported by Liang *et al*.. **(B)** Number of viral species identified by Phanta in all metagenome pairs, from both infant cohorts. Each dot represents a metagenome and lines connect metagenome pairs. **(C)** Overlap between viral species identified by Phanta in 10 pairs of metagenomes (see Methods) from infants in the four-month cohort. Each bar represents the total relative taxonomic abundance of viruses identified in a virus-enriched metagenome. Colors depict the proportion of this abundance from species also found in the paired bulk metagenome. **(D)** Complementary analysis to (C), showing the proportion of relative taxonomic abundance in bulk metagenomes from species also found in virus-enriched metagenomes. **(E)** Abundance ratio of virulent to temperate species detected by Phanta in virus-enriched and bulk metagenomes from the four-month cohort. Ratios were obtained using one of Phanta’s provided post-processing scripts, along with viral lifestyle predictions that were made by BACPHLIP and are provided in Phanta’s default database. **(F)** Phanta’s abundance estimates for *Bifidobacterium* and predicted *Bifidobacterium* phages in bulk metagenomes from infants in the four-month cohort (who had a range of diets). This analysis was facilitated by one of Phanta’s provided post-processing scripts, along with host genus predictions that were made by iPHoP^65^ and are provided in Phanta’s default database. **(G)** Relationship between the originally reported abundance of Adenovirus in infant stool samples (based on qPCR), vs. the newly determined abundance, based on applying Phanta to the corresponding metagenomes. This analysis considered all stool samples from the four-month cohort; most were negative or weakly positive by both methods (i.e. plotted close to (0, 0)). **(H)** Heatmap of Adenovirus abundance in stool samples from infants in the four-month cohort, as determined by four complementary methods. Shown are stool samples originally reported to be positive for Adenovirus by qPCR. Method abbreviations: qPCR = qPCR for Adenovirus from DNA extracted from virus-like particles; quantified by genome copies / μl DNA. Assembly = alignment of assembled contigs from bulk metagenomes to Adenovirus genomes; quantified by number of contigs identified as Adenovirus. Bulk/virus-enriched = application of Phanta to bulk or virus-enriched metagenomes, using the default Phanta database; quantified by genome copies per million reads. **Note:** in all boxplots, boxes represent the interquartile range (IQR), the horizontal line indicates the median, and whiskers extend between (25th percentile - 1.5*IQR) and (75th percentile + 1.5*IQR).

### Viral profiles from bulk and virus-enriched metagenomes overlap, but complement each other

Given its high sensitivity, we hypothesized that Phanta would detect a comparable number of viral species in bulk metagenomes as in virus-enriched metagenomes. Indeed, the number of species detected was similar in paired bulk and virus-enriched metagenomes (Fig. 5B). We further tested whether the bulk and viral-enriched metagenomes provide a similar profile of the viral community by examining pairs of bulk and viral-enriched metagenomes from the same sample. First, we examined 10 pairs of metagenomes with relatively deep sequencing of the bulk metagenomes (range of 150bp paired-end reads 8.6M - 13.3M; median = 9.3M). Species present in bulk metagenomes captured a median of 94% of the viral abundance in virus-enriched metagenomes (Fig. 5C). The variance in this quantity is mostly explained by the sequencing depth of the bulk metagenomes (Supplementary Fig. 5). To complement this analysis, we examined 10 pairs of metagenomes with highly successful viral enrichment (see Methods). Species present in virus-enriched metagenomes captured a median of 69% of the viral abundance in bulk metagenomes (Fig. 5D). Those differences are expected as shotgun metagenomes can capture viruses that did not enrich in the VLP enrichment process for a variety of reasons, technical or biological^31^. For example, prophages lack viral-like particles, and are therefore more likely to be captured by bulk metagenomes. Given the inclusive nature of bulk metagenomes, they capture more viral species per total number of viral reads (Supplementary Fig. 6A), whereas viral-enriched metagenomes capture more viral species per total number of metagenome reads (Supplementary Fig. 6B). With the ability to identify prophages in bulk metagenomes, we hypothesized that the fraction of temperate phages would be higher in virus-enriched metagenomes. Indeed, we observed a 3-fold higher virulent/temperate abundance ratio in virus-enriched metagenomes relative to bulk (Fig. 5E; Supplementary Fig. 6C).

### Phanta is highly effective for simultaneous quantification of phages and their hosts from a single metagenomics experiment

One advantage of using Phanta to profile bulk metagenomes, as opposed to virus-enriched metagenomes, is the ability to examine phages and their hosts simultaneously and from a single dataset, instead of two separately generated datasets. Using a Phanta-based analysis of the bulk metagenomic dataset from Liang *et al*.^59^ investigating the impact of diet on the infant gut, we found that Bifidobacterium and its phages are ~2-fold more abundant in breastfed infants relative to formula-fed or infants that were fed with a mixed breast milk and formula diet (Fig. 5F). This observation, although expected, demonstrates the power of Phanta to simultaneously identify phages and their bacterial hosts and to associate them with known traits.

### Phanta accurately identifies and quantifies human-infecting viruses

Lastly, we wished to test the ability of Phanta to accurately identify human-infecting viruses in metagenomes. Liang *et al*. were able to identify viruses in the family of Adenoviridae using qPCR from their infant stool samples^59^. Phanta identified 5 samples with the mastadenovirus C species, with almost perfect correlation between the estimation of genome copies per uL using qPCR and Phanta’s estimation of genome copies per million reads (Fig. 5G). Phanta was able to identify Adenoviruses in bulk shotgun metagenomic samples with as low as 88 copies/uL in qPCR and successfully identified all samples with >550 copies/uL. Phanta demonstrated higher sensitivity in identifying Adenoviruses relative to using assembly-based methods (Fig. 5H), which only detected Adenoviruses in samples that had >20,000 copies/uL by qPCR. Of note, we used the assembled contigs to confirm that Phanta successfully identified the right Adenovirus species, by aligning the contigs to all Adenovirus genomes from RefSeq.

## Discussion

A major goal of microbiome studies is to identify microbial features associated with traits of interest, such as phenotypes, lifestyle factors, and health status. In an ideal world, organisms from all domains could be accurately quantified in a single experiment. The first step in achieving this goal is to profile microbial communities - i.e., to determine their composition from sequencing data. Although shotgun metagenomes capture both prokaryotes and viruses, profiling the viral fraction of microbial communities has historically presented a greater challenge and has required specially tailored methods. For example, popular reference-based methods have allowed accurate profiling of prokaryotes from metagenomes^24^ without being able to accurately capture viruses due to the historical lack of comprehensive reference databases of viral genomes^27^.

Because of these limitations, profiling viruses has required additional orthogonal analyses, based on assembling metagenomic reads and identifying viral genomes *de novo*^25,26^. In addition, due to the relatively low abundance of viral sequences in bulk metagenomes, it has been common to conduct an entirely separate experiment to profile the virome by enriching for viral sequences prior to making sequencing libraries^58^.

With the recent development of much more comprehensive databases of viral genomes^27,31–34^, deeply sequenced bulk metagenomes, and fast and accurate read classifiers^22,38^, technical and experimental advances have converged to make it possible to integrate prokaryotic and viral profiling. By harnessing the latest developments, Phanta enables simultaneous profiling of bacteriophages and their prokaryotic hosts, in a single experiment and with a single analysis. This simultaneous profiling has several advantages. First, it reduces the need to sequence both viral-enriched and bulk metagenomes, thus reducing research time and costs, in addition to eliminating technical differences between two separate experiments. Second, it bypasses the need to use separate computational workflows to profile prokaryotes and viruses. Lastly, and most importantly, it allows the study of cross-domain interactions between phages and their hosts, either in novel datasets, or in the wealth of metagenomic datasets that are already publicly available.

Although Phanta can be applied with different databases, Phanta’s default database was constructed with the human gut in mind. For decades, the viral portion of the human gut was mostly unknown, and considered as “dark matter”^60,61^. There is still much to learn, with some basic discoveries occurring only in the past few years. For example, the first representative of one of the most abundant bacteriophage clades - crAss-like viruses - was discovered only in 2014^56^. Similar examples, such as the highly prevalent Hankyphage (p00)^57^ and LoVEphage^37^, were discovered only in 2018 and 2021, respectively. We anticipate that Phanta, when applied with its default database, will allow similar key discoveries to be made. In this study alone, we were able to estimate a ~2:1 ratio of viruses to bacteria in the human gut, determine that temperate and Bacteroides-infecting phages dominate the gut phageome, and demonstrate a high interindividual variability of the gut virome, as compared to the bacteriome. These and other core principles can serve as a springboard for more extensive discovery, such that “gut microbiome” will no longer be publicly synonymous with “gut bacteria,” but rather understood as a complex community with many types of interacting members.

Importantly, Phanta was developed with careful attention to the risk of spurious discovery, as read-based classifiers are frequently known to make mistakes, and thus to identify false positive taxa^40^. As described, to mitigate false classification we increase classification confidence and filter out species with low genome coverage, an idea that was previously described in the implementation of KrakenUniq^40^. Of course, these decisions come with potential costs. For example, increasing the required confidence of classification may lead reads from some species to all classify at higher taxonomic ranks during the Kraken2 step of the workflow. In such a scenario, the sensitivity of viral identification would be decreased, since during Bracken, classified reads are only redistributed to species that initially received some direct classifications. Similarly, requiring a certain genome coverage reduces the probability of identifying lowly abundant species with long genomes. However, all the relevant parameters of Phanta are user-adjustable, and using our simulations we were able to show that a combination of minor increments in both thresholds is sufficient to reduce most of the noise with a very small cost to signal (Figs. 2D-E).

More broadly, Phanta offers a flexible setup that can be modified according to the user’s analysis goals and main concerns. If a user aims to minimize false negatives, i.e. to increase the probability of identifying all species while allowing a substantial increase in false positives, the user can decrease (1) the confidence cutoff, (2) the coverage requirement, and (3) the minimal number of reads directly classified to a species for it to receive an abundance estimate. On the other hand, if a user wishes to minimize false positives while taking the risk of decreasing true positives, the user can increase these three parameters. Phanta also provides an alternative database to the default, in which predicted prophages in the HumGut genomes were masked. This masked database can be used to increase the likelihood of identifying prophages. In addition to parameters and database choice, the characteristics of a sequencing experiment can impact the power of identification by Phanta. Although we did find high agreement between viral-enriched and bulk shotgun metagenomes (Fig. 5C), enriching the library for viral particles would be recommended if a researcher (i) prioritizes identification of viruses that are particles over prophages, (ii) is not focused on determining cross-domain interactions, and (iii) is limited by the possible depth of metagenomic sequencing. Conversely, bulk metagenomic analysis allows users to: (i) profile prophages in addition to virulent phages, (ii) avoid potential biases introduced by the process of isolating viral particles, and (iii) identify cross-domain interactions, both within and across samples. Given the low and rapidly decreasing costs of shotgun sequencing, and our findings that bulk metagenomes of fairly standard depth allow for comparable virus identification to viral particle-enriched fractions, we anticipate that many researchers may opt to enhance their standard analyses of bulk metagenomes by applying Phanta.

While Phanta enhances the knowledge that can be gained about viruses from bulk metagenomes, it has several limitations. First, while Phanta has high sensitivity, using a reference-based method restricts identifications to the genomes in the database, and thus limits resolution. For example, Phanta’s default database is biased toward dsDNA viruses identified in the human gut. Similarly, while Phanta does include some eukaryotes in its default database, our knowledge of this domain in the gut is still limited; this is, in part, due to limitations in reference databases for protists, amoeba, helminths and fungi. Improvements in eukaryotic reference databases should enhance eukaryote classification in the coming years. Second, extending Phanta to characterize the virome in other human microbiomes, such as the skin or vaginal microbiome, may require curation of additional metagenome-derived virome databases generated from these niches. Furthermore, classifying short reads to reference genomes is challenging when reads originate from genomic regions that are conserved between species. Moreover, the usage of *k*-mer-based methods, although fast and computationally efficient, does not provide information required for aligning reads to a specific region in the genome, and thus does not allow investigation of genome variation. Finally, viral taxonomy is not as well-defined as prokaryotic taxonomy, and thus Phanta cannot currently provide specific named designations to many viral species, beyond family- or order-level assignments. We anticipate that as knowledge of the virome increases, this challenge will begin to be addressed.

Despite these limitations, Phanta is benchmarked, easy to use, carefully tuned to limit false positives, and able to provide simultaneous profiling of various domains from a single experiment. These advantages suggest that Phanta will help accelerate the study of the virome in human gut microbiomes, as well as illuminate cross-domain interactions in this niche. Phanta enables much higher resolution of the viral portion in a human gut sample when analyzing a bulk metagenome relative to current approaches or databases, and thus it promises to provide exciting insights when applied to the tens of thousands of human gut metagenomes that have already been sequenced, to date. We expect that Phanta will be both: (1) used to re-analyze publicly available data, and (2) taken into account when planning new experiments. Overall, Phanta lowers the barrier to virus-inclusive studies of the gut microbiome, and we expect that its application will confidently identify numerous novel associations between viruses, prokaryotes, and human traits.

## Online Methods

### Constructing a comprehensive, taxonomy-aware, domain-inclusive database of human gut microbes

Phanta’s default database was constructed to be compatible with the Kraken2/Bracken tools^22,38^. Therefore, its construction required curating: 1) a large collection of genomes, and 2) taxonomy files placing each genome within a tree of named nodes.

The viral genomes within the database were sourced from: 1) the recently published human gut-focused MGV catalog (available at (https://portal.nersc.gov/MGV/)^27^ and 2) RefSeq^21^, the database of reference genomes maintained by NCBI (MM/YY of download: 02/22).

After downloading the viral genomes, the viral taxonomy tree was constructed. The first step was to download the complete NCBI taxonomy using the kraken2-build --download-taxonomy utility. Next, branches of the taxonomy were pruned so that only the branches leading to the RefSeq viral genomes remained.

After providing taxonomic assignments to RefSeq genomes, assignments were provided to the MGV genomes. The first step in doing so was to group the MGV genomes into the 54,118 ANI-based species specified by the MGV paper^27^. Each of these species came with a designated “species representative genome” that was chosen based on features such as circularity and length. Code on the MGV GitHub page (https://github.com/snayfach/MGV/tree/master/aai_cluster) was then used to cluster species into genera based on amino acid identity (AAI) and gene sharing between the representative genomes.

To avoid species duplications between MGV and RefSeq viruses, and to provide a full NCBI taxonomy for MGV genomes where available, average nucleotide identity was calculated between all of the 54,118 species representative genomes in MGV and all the RefSeq viral genomes using fastANI^62^. In cases where an MGV species representative genome had > 95% ANI to a RefSeq viral genome, all of the genomes in the relevant MGV species were re-assigned to RefSeq, i.e., designated as strains of the RefSeq viral genome.

To determine where each AAI-based MGV genus fit into the NCBI taxonomy, we utilized a file from the MGV website (https://portal.nersc.gov/MGV/) that provides - when possible - NCBI-recognized taxonomic annotations for each genome at the genus, family, and/or order levels, based on amino acid alignments to a protein database^27^. We used this information to remove some of the AAI-based genera and re-assign their contained species to the relevant NCBI-recognized genus. Specifically, for each AAI-based genus, we calculated the percentage of species representative genomes within the genus that had an NCBI genus annotation provided. If this percentage was greater than 50%, and the NCBI genus annotation was consistent for > 90% of the species representative genomes with annotations, the AAI-based genus was removed and all of its species were re-assigned to the NCBI genus.

The remaining AAI-based genera were then assigned as direct descendants of the lowest possible NCBI-recognized taxonomic level, by iterating a variant of the strategy described above. More specifically, starting with family: if > 50% of the species representative genomes within a given AAI-based genus had an NCBI family annotation, and the NCBI family annotation was consistent for > 90% of the species representative genomes with annotations, the AAI-based genus was assigned as a direct descendant of the relevant NCBI family. The remaining AAI-based genera (i.e., those without a family assignment) were then assigned to an order - when possible - in the same manner. All of the AAI-based genera without an order assignment were assigned as direct descendants of the superkingdom of Viruses.

The prokaryotic genomes within Phanta’s database were sourced from HumGut, a recently published human gut-focused catalog of prevalent bacterial and archaeal genomes^19^. The HumGut catalog was in turn sourced from both the Unified Human Gastrointestinal Genome (UHGG) collection^18^ and RefSeq^21^. An NCBI-compatible taxonomy file for the HumGut genomes was downloaded directly from the HumGut website (http://arken.nmbu.no/~larssn/humgut/). The branches of the NCBI taxonomy leading to the human genome were also included in this taxonomy file and thus we also included the human genome (hg38) in our database.

We sourced fungal genomes from RefSeq and common contaminant sequences from the Core UniVec database using the kraken2-build download-library command provided by the Kraken2 developers (MM/YY of download: 02/22). The relevant branches of the NCBI taxonomy were then obtained in the same way that they were obtained for the RefSeq viral genomes (i.e., by “pruning” the full NCBI taxonomy, please see above).

Finally, the constructed taxonomy files for each portion of the database were concatenated, and a Kraken2/Bracken-compatible database was built using the commands provided on the Github sites (https://github.com/DerrickWood/kraken2; https://github.com/jenniferlu717/Bracken).

### Masking prophages in prokaryotic genomes

An alternative version of Phanta’s default database was also created, in which predicted prophages were masked (i.e., replaced with Ns) within all the prokaryotic genomes from HumGut. VIBRANT (v1.2.1)^42^ was used to predict prophages within the HumGut genomes. Prophage coordinates were extracted and masking was conducted using the bedtools utility MaskFastaFromBed^63^. All of the analyses in this paper were conducted using the unmasked version of the database, except where explicitly noted otherwise.

### Workflow implementation

Phanta was implemented using the workflow management system Snakemake. Core scripts are written in Python, bash, and R. A step-by-step tutorial detailing workflow installation and usage is provided on the main page of the Phanta GitHub (https://github.com/bhattlab/phanta). Briefly, after cloning the GitHub repository to their system, users should: 1) download the desired database - default (unmasked) or masked - via the command line, 2) make slight edits to a configuration file, and 3) execute the provided Snakemake command on the command line, within the appropriate conda environment that is fully specified in a provided yaml file. As detailed in the GitHub tutorial, the repository also provides a test data set that can be used to verify that the workflow was installed correctly.

### Classification of metagenomic reads to taxa

The first step of the Phanta workflow is classification of metagenomic reads in each sample against the desired database of genomes (default/unmasked or masked, see above). Classification is conducted using the Kraken2 tool (currently v2.1.2)^22^, which classifies reads using a *k*-mer-based approach. More specifically, to classify each read, Kraken2 slides along the read length, computes a “minimizer” (i.e., compact version) of each *k*-mer, and looks up where the minimizer maps in the genome database. After all the minimizers in the read have been looked up, Kraken2 classifies the read to the lowest taxonomic level possible, considering the user’s preference for the confidence in the assignment (supplied via the --confidence parameter to Kraken2). By default, Phanta supplies a confidence of 0.1 to Kraken2, but this value can be adjusted by the user in the Snakemake configuration file. This parameter ranges from 0 to 1 and essentially specifies a certain fraction of a read’s *k*-mers to be mapped to a given taxon, in order for Kraken2 to make that classification. E.g., 0.1 = 10%.

Phanta also makes use of the --report-minimizer-data parameter available in Kraken2 v2.1.2, that is based on ideas from KrakenUniq^40^. Providing this parameter modifies the standard Kraken2 output to report an additional data point for each taxon, specifically: how many unique minimizers in the genomes of this taxon are covered by read sequences?

### Filtering of false positive species after classification

Phanta filters likely false positive species from each sample after the initial classification step and before species-level abundance estimates are calculated (Figure 1). This filtering step makes use of the minimizer data reported by Kraken2 during the classification step (described above, in the section “Classification of metagenomic reads to taxa”).

Specifically, a proxy for genome coverage is calculated for each genome of each species identified during classification. This proxy is calculated by dividing: 1) the reported number of unique minimizers in the genome that are covered by read sequences, by 2) the total number of unique minimizers contained in the genome. The denominator of this fraction is not reported in the Kraken2 output, but is obtained by Phanta from an “inspect.out” file contained within the genome database (originally generated using the kraken2-inspect functionality).

Bacterial and viral species are marked as false positives and filtered out if none of their strain-level genomes have a calculated coverage above a user-specified threshold. Suggested thresholds are provided in the Snakemake configuration file (0.01 for bacterial species; 0.1 for viral species). These suggested thresholds were chosen because they yielded a high signal-to-noise ratio in identified species when tested on the mixed simulated metagenomes described below.

Users can also require that the numerator of the fraction above (i.e., the number of unique minimizers covered by reads) be higher than a specified threshold for at least one strain-level genome of each “true positive” species. In other words, it is possible to specify that a high calculated genome coverage will not “count” unless the number of unique minimizers is higher than a specific value (e.g., > 300 unique minimizers) for at least one strain-level genome. By default, this option is not utilized by Phanta but can be implemented by the user by making use of the minimizer_thresh_viral and minimizer_thresh_bacterial parameters in the Snakemake configuration file.

### Species abundance estimation and correction for genome length

After species are filtered from the Kraken2 output, abundances of the remaining species are estimated using the Kraken2-compatible tool Bracken (currently v2.7)^38^. Bracken estimates species-level abundances by redistributing all classified reads to the species level.

Of note, Bracken accepts a threshold parameter that specifies one last filter for false positive species - how many sample reads must have been classified to a species during Kraken2 classification for Bracken to estimate its abundance? By default, Phanta specifies this threshold as 10 reads - the accepted standard for running Bracken - but this number can be adjusted by the user through the filter_thresh argument in the Snakemake configuration file.

We also utilize Bracken output to calculate relative taxonomic abundance estimates for each species by considering genome length. Specifically, the abundance estimate for each species is scaled by the median length of the genomes under the species. Additional normalizations are also provided in this corrected output file, such as reads per million reads per million base pairs (analogous to RPKM in transcriptomics), copies per million reads, and more.

### Provided post-processing scripts

There are three main post-processing scripts in the Phanta GitHub.

The first calculates “lifestyle statistics” for the viral community in each metagenome (e.g., ratio of virulent:temperate viruses), based on lifestyle predictions for viral species that are provided in Phanta’s default database. Lifestyle predictions for species from MGV were obtained from the mgv_contig_info file provided in the MGV database^27^. These predictions were calculated using BACPHLIP^64^ and we used the same tool (v0.9.6) to make lifestyle predictions for viral species from RefSeq. Throughout the manuscript, viruses with a BACPHLIP-predicted virulence score above 0.5 were considered virulent; others were considered temperate.

The second collapses viral abundances in each sample by predicted host, based on provided host predictions for viral species in Phanta’s default database. Host predictions were made using iPHoP^65^.

The third correlates the abundances of bacterial and viral species in each sample. This cross-kingdom correlation is done by fastspar^66,67^- a method designed to correlate compositional data.

Also provided are post-processing scripts to filter or sum abundance tables (counts, relative read abundances, or relative taxonomic abundances) to a desired taxonomic rank (e.g., species or genus).

### Simulating mixed metagenomes

10 mixed metagenomes (each containing ~6.5M paired-end 150bp reads) were simulated using CAMISIM (v1.3)^68^. These simulated metagenomes were used to generate the data in Figure 2. Each simulated metagenome consisted of: 1) 95% prokaryotic reads from 300 randomly chosen genomes from the HumGut catalog, and 2) 5% viral reads from 50 randomly chosen genomes from the MGV catalog.

### Download and processing of publicly available, short-read human gut metagenomes

245 shotgun gut metagenomes from healthy human adults in a Japanese cohort were downloaded from SRA (accession DRP004793 - Yachida *et al*.^43^). Shotgun gut metagenomes from infants were also downloaded from SRA (accession PRJNA524703 - Liang *et al*.^59^). The full list of downloaded samples, along with accession numbers, is available within Supplementary Table 2.

Following download, each metagenome was preprocessed as follows. First, reads that exactly matched each other (PCR duplicates) were removed using hts_SuperDeduper (v1.2.0). Next, TrimGalore (v0.6.5 healthy adults; v0.6.7 infants) was used to: 1) trim low-quality bases (Phred score < 30) from the ends of reads, and 2) discard reads with a final length of < 60bp. Human reads were then removed using BWA alignment against the human genome (GRCh37). Initial and final quality checks were performed using MultiQC (v1.7 healthy adults; v1.11 infants).

All results from applying Phanta to these metagenomes were obtained using Phanta’s default database and parameters, except where explicitly noted otherwise (i.e., varied databases were tested in Figures 3A and 3B). Note also that for the infant cohort, the database file required for running Bracken was slightly modified from default (adjusted for 120bp reads rather than 150bp, following the instructions on the Bracken GitHub).

Separate from running Phanta, a subset of these metagenomes was assembled into contigs and scaffolds using metaSPADES^50^ version 3.15. Specifically, the following metagenomes were assembled: 1) fifty randomly selected metagenomes from the healthy adult cohort, and 2) all bulk metagenomes from the “four-month” subgroup of the infant cohort. The specific metagenomes that were successfully assembled are indicated in Supplementary Table 2.

### Assembly-based method for identifying viral species in healthy adult gut metagenomes

To generate the results in Fig. 3C, the 50 assembled healthy adult gut metagenomes were run through two standard methods for phage identification from metagenomic assemblies. The first method, VIBRANT, uses a hybrid machine learning and protein similarity approach to identify viral signatures^42^. The second method, VirSorter, predicts protein-coding genes in assembled DNA sequences and assesses their similarity to known viral proteins^45^.

VIBRANT (v.1.2.0) was run on assembled scaffolds and the quality and completeness of identified phages were estimated by CheckV (v.0.7.0)^51^ using database v0.6. Low-quality phage scaffolds were filtered out.

A similar procedure was performed using VirSorter (v1.0.6, downloaded in February 2018), where phage contigs were classified as category 1, 2, or 3 depending on confidence level. Category 3 predictions were filtered out before running CheckV.

Finally, dRep (v3.2.2)^69^ was applied to the combined set of quality-filtered phage contigs predicted by VIBRANT+VirSorter in each sample to extract a unique set of phage genomes based on an ANI threshold of 0.95 and coverage threshold of 0.5. fastANI was applied for secondary clustering and genome filters included a minimum length of 1000 bp, an N50 weight of 0, and a size weight of 1.

The full list of parameters utilized with the “drep dereplicate” utility was: *-sa 0.95 --S_algorithm fastANI -nc .5 -l 1000 -N50W 0 -sizeW 1 --ignoreGenomeQuality --clusterAlg single*

### Assembly-based method for identifying Adenoviruses in stool samples

The assembled bulk metagenomes from the “four-month” subgroup of the infant cohort were used to calculate the column labeled “Assembly” in Fig. 5H. FastANI was used to calculate average nucleotide identity between all assembled contigs in each sample and 1801 *Adenoviridae* genomes available in NCBI (retrieved by *datasets download genome taxon Adenoviridae*). Each contig with ANI score >=95% to at least one *Adenoviridae* genome was counted as an Adenovirus.

### Calculation of dissimilarities between metagenomes

Bray-Curtis and Jaccard distances were calculated using the R package vegan, version 2.5-7.

### Choosing pairs of metagenomes for overlap analysis

For the analyses in Figs. 5C-5D, we wanted to determine how well each type of metagenome could represent the information in the other, excluding samples with low sequencing depth of the bulk metagenomes, or low enrichment of virus-enriched metagenomes.

For the analysis in Fig. 5C, we chose pairs of metagenomes with decent viral enrichment and deeply sequenced bulk metagenomes. Specifically: (1) We identified the top 50% of samples based on the percent of reads that Phanta assigned to viruses in the virus-enriched metagenomes; (2) Of these, we selected 10 samples whose paired bulk metagenomes were the most deeply sequenced..

For the analysis in Fig. 5D, we chose pairs of metagenomes with decent bulk sequencing depth and highly successful viral enrichment. Specifically: (1) We identified the top 50% of samples based on the sequencing depth of the bulk metagenomes; (2) Of these, we selected 10 samples with the highest percent of reads that Phanta assigned to viruses in the virus-enriched metagenomes.

### Determination of size and number of genomes in each Kraken2/Bracken-compatible database tested

To determine the size of each Kraken2/Bracken-compatible database tested (listed in Table 1), we summed the sizes of the following files and rounded to the nearest GB: hash.k2d, opts.k2d, taxo.k2d, seqid2taxid.map, database150mers.kmer_distrib. These are the files necessary for running Kraken2 and Bracken. We obtained the number of prokaryotic and viral genomes in each database that we did not construct directly from the relevant publications: Wright *et al*., 2022 (for Standard Kraken2 and RefSeq Complete)^39^; Almeida *et al*., 2021 (for UHGG)^18^; Hiseni *et al*., 2021 (for HumGut)^19^.

## Data and code availability

Phanta is publicly available at https://github.com/bhattlab/phanta with a detailed tutorial describing installation and usage. Accession numbers of all publicly available metagenomes used for analysis are provided in Supplementary Table 2. Workflows used for preprocessing and assembly were used in this manuscript and are available at: https://github.com/bhattlab/bhattlab_workflows.

## Acknowledgements

We thank Dylan Maghini and Boryana Doyle for thoughtful comments on the manuscript, Stephen Nayfach and Pranvera Hiseni for helpful conversations, Jakob Wirbel for testing Phanta, and Ben Siranosian and the Stanford Research Computing Center for computational support. Computing costs were supported, in part, by a NIH S10 Shared Instrumentation Grant 1S10OD02014101. Figure 1 was created using BioRender.com. This work was supported in part by NIH R01AI14862302 & R01AI14375702, a Stand Up 2 Cancer Grant, the Chan Zuckerberg Initiative, a Sloan Foundation Fellowship and the Allen Distinguished Investigator Award (to A.S.B.). Y.P. is supported by the School of Medicine Dean’s Postdoctoral Fellowship. M.C. is supported by an NIH-funded predoctoral fellowship (5T32HG000044-25) and the National Defense Science and Engineering Graduate Fellowship (starting September 2022).

