## Supplementary Figures for "Phanta: Phage-inclusive profiling of human gut metagenomes"

1. Department of Genetics, Stanford University, Stanford, CA, USA.
2. Department of Medicine, Divisions of Hematology and Blood & Marrow Transplantation, Stanford University, Stanford, CA, USA.
* These authors contributed equally

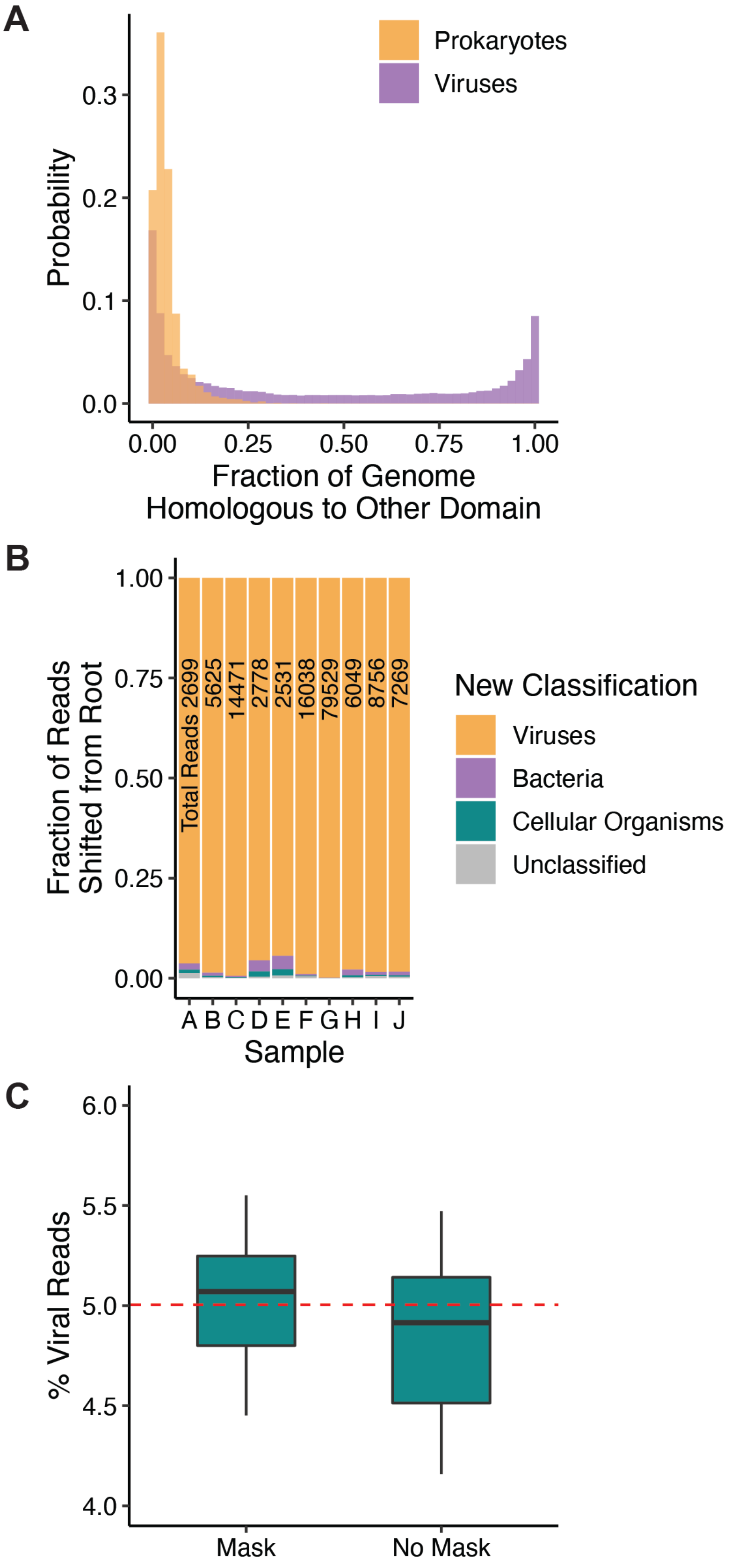


**Supplementary Figure 1. Effect of masking prophages within prokaryotic genomes in Phanta's default database.** **(A)** Probability distribution of the fraction of viral and prokaryotic genomes that are homologous to the other domain. Homologous regions were retrieved by aligning all viral genomes from the MGV catalog[27] and prokaryotic genomes from the UHGG collection[18]. **(B)** Post-masking classification of reads that were originally classified to the "root" node of the taxonomy (i.e., ambiguous domain) but received a new classification after masking. Bars represent the 10 simulated metagenomes from Figure 2. The number in each bar indicates the total number of reads that the bar represents. **(C)** Percent of reads in simulated metagenomes assigned to viruses with and without masking. Dashed red line depicts the true percentage of viral reads.

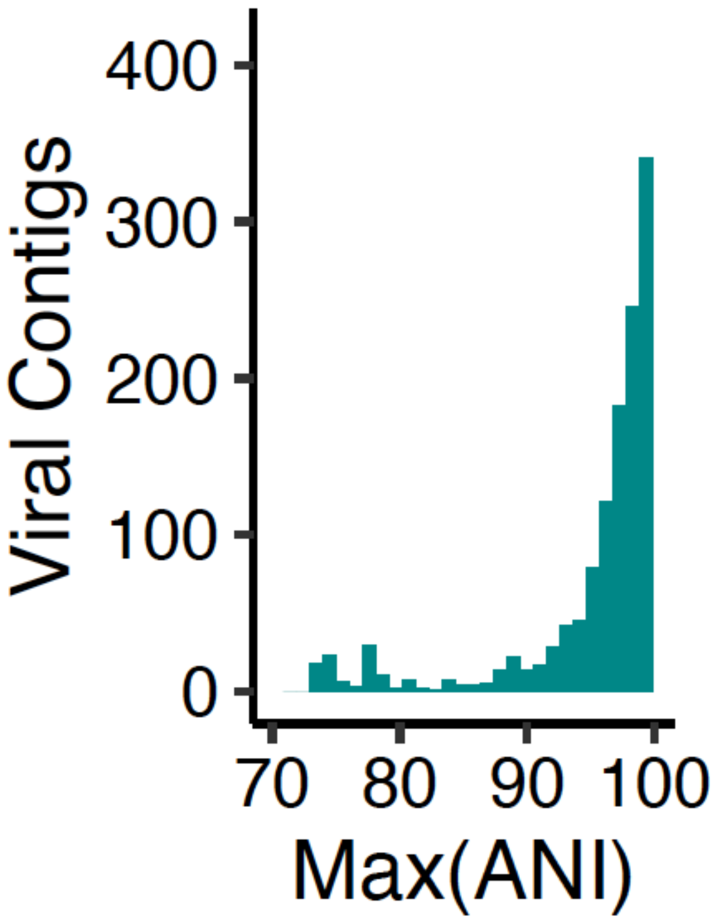


**Supplementary Figure 2. Identity between phages identified by a standard assembly-based method and phages identified by Phanta in healthy adult metagenomes.** Distribution of maximum ANI score between each contig identified in the assembly-based workflow from Fig. 3C and the set of viral genomes in Phanta’s default database.

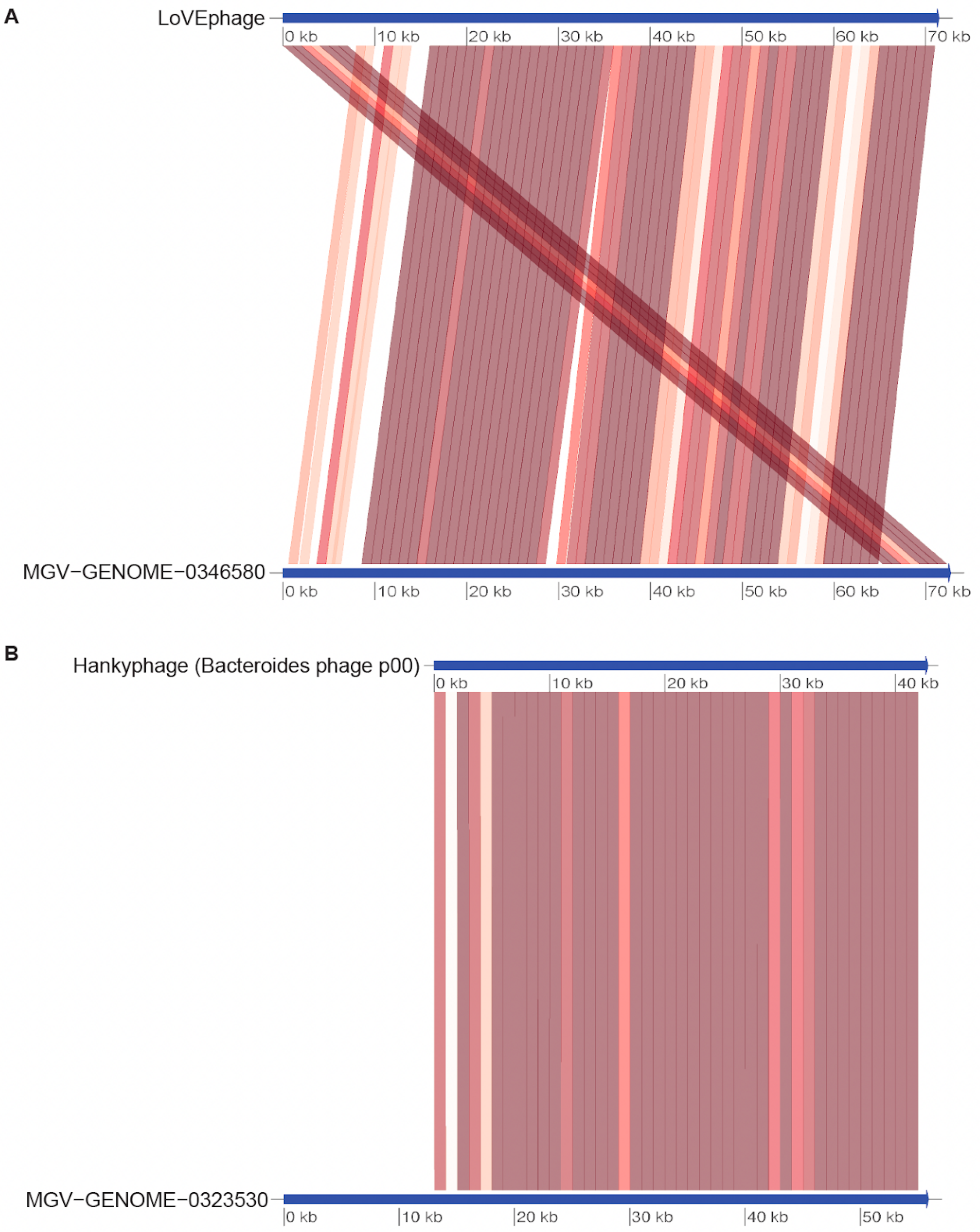


**Supplementary Figure 3.** Genome-wide average nucleotide identity (ANI) between the species representative genomes of Caudovirales OTUs 66229 and 72541 (in Phanta's default database), and the genomes of their high-confidence BLAST hits, which are:
**(A)** LoVEphage[37] (GenBank: MZ919987.1), and
**(B)** Hankyphage (Bacteroides phage p00; GenBank: BK010646.1)[56], respectively. Genome-wide ANI was 99.9% and 98.9%, respectively, as determined by fastANI. Each connection between genomes represents a bidirectional mapping between 1000bp fragments. Colors represent ANI [90-100%]; darker colors represent higher ANI.

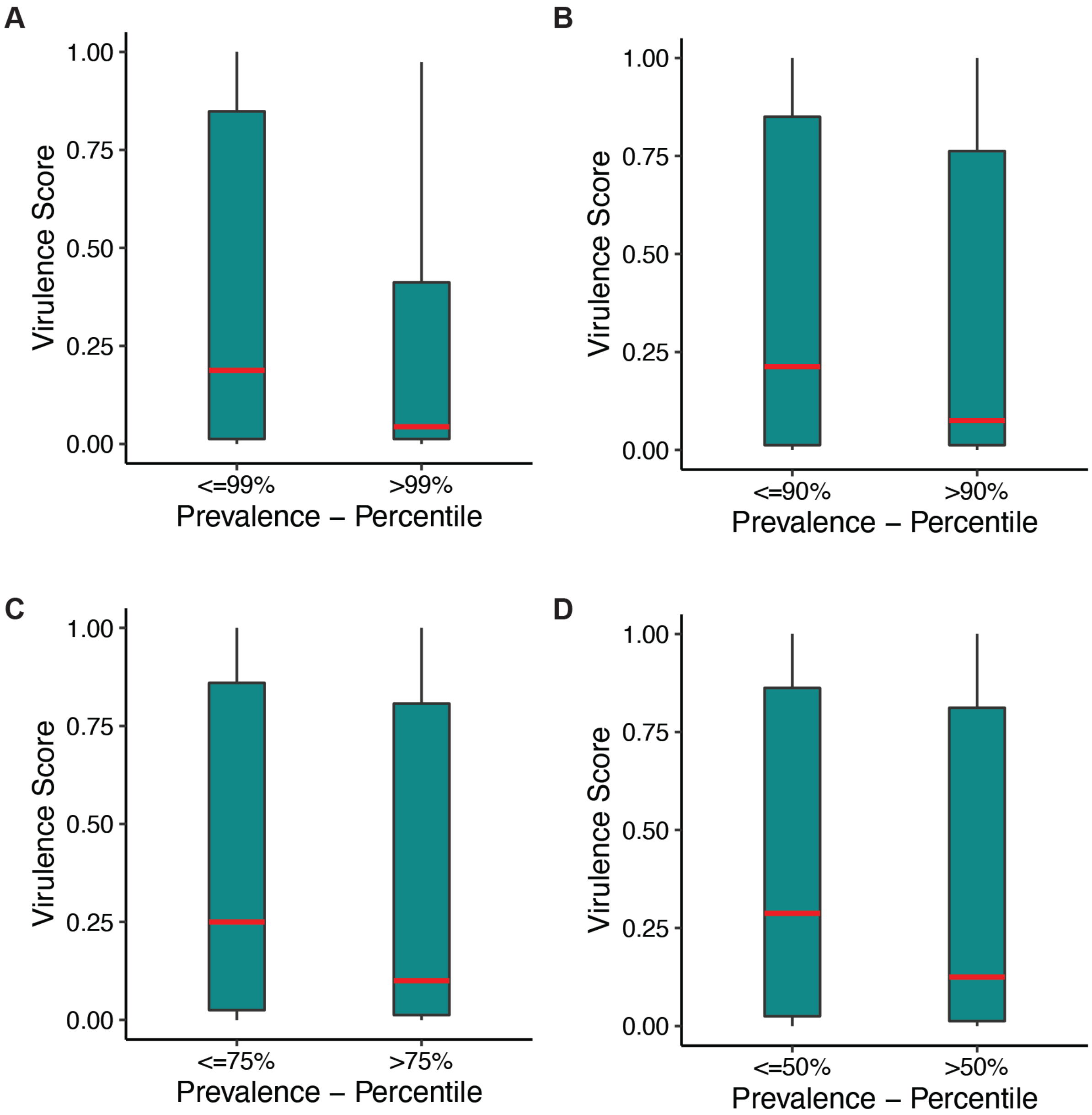


**Supplementary Figure 4. Lifestyle of prevalent phages.** Comparison of virulence score (predicted by BACPHLIP) between the more prevalent viral species in the healthy adult cohort, vs. the other viral species. Each panel shows a different cutoff for defining "more prevalent species" - e.g., panel **(A)** compares the top 1% of species in terms of prevalence, vs. the bottom 99%. Similarly, panel **(B)** - **(D)** compare: (B) the top 10% vs. the bottom 90%, (C) the top 25% vs. the bottom 75%, (and (D) the top 50% vs. the bottom 50%.

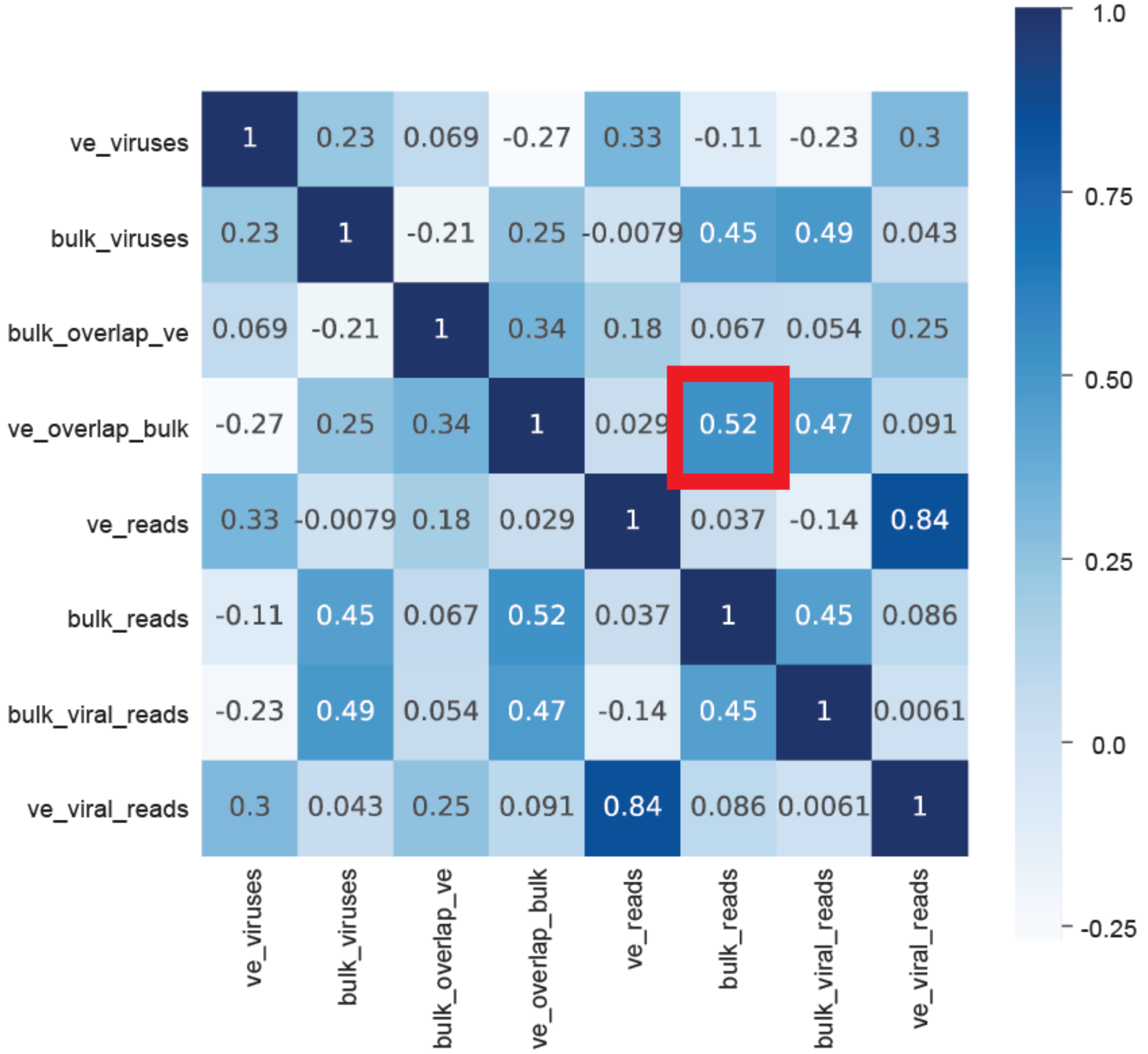


**Supplementary Figure 5. Correlations between statistics related to viral content in bulk and virus-enriched gut metagenomes from the infant gut.** All-by-all correlations between statistics related to viral content in shotgun gut metagenomes from the four-month cohort of Liang *et al.*. All of these statistics were calculated in this study and most are based on outputs of the Phanta pipeline. Specifically, eight statistics were correlated: ***(1) ve_viruses:*** the number of viral species in virus-enriched metagenomes; **(2)** ***bulk_viruses:*** same as (1) but for bulk metagenomes; ***(3) bulk_overlap_ve:*** the proportion of the viral abundance in the bulk metagenome from species that were also detected in the virus-enriched metagenome; ***(4) ve_overlap_bulk:*** same as (3) but reversed; ***(5) ve_reads:*** the total number of reads in the virus-enriched metagenome; ***(6) bulk_reads:*** the total number of reads in the bulk metagenome; ***(7) bulk_viral_reads:*** the total number of reads assigned to viruses in the bulk metagenome; ***(8) ve_viral_reads:*** the total number of reads assigned to viruses in the virus-enriched metagenome. The highlighted box shows the correlation between the sequencing depth of the bulk metagenome and ***ve_overlap_bulk***, i.e. the proportion of viral abundance in the virus-enriched metagenome from species that were also detected in the shotgun metagenome.

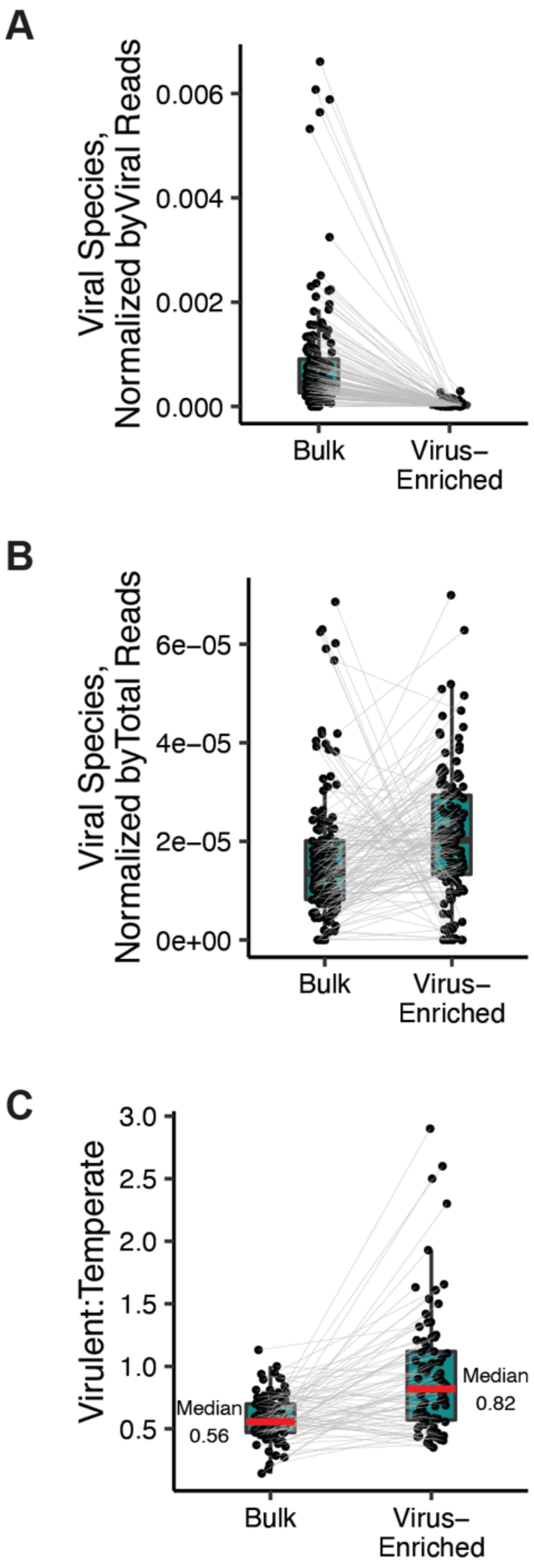


**Supplementary Figure 6. Application of Phanta to virus-enriched and bulk shotgun metagenomes from the infant gut.**
**(A)** Number of viral species identified by Phanta - normalized by total reads assigned to viruses - in paired virus-enriched and bulk shotgun metagenomes from infants in both infant cohorts (longitudinal and four-month) from Liang *et al.*. Each dot represents a metagenome and lines connect paired metagenomes (i.e., those from the same original stool collection).
**(B)** Same as (A) but normalized by total number of reads in the metagenome.
**(C)** Ratio of number of virulent viral species to number of temperate viral species detected by Phanta in virus-enriched vs. bulk shotgun metagenomes from infants in the four-month cohort. Ratios were obtained using one of Phanta's provided post-processing scripts.
